# Patterns of compensatory mutations in *rpoA/B/C* genes of multidrug resistant *M. tuberculosis* in Uganda

**DOI:** 10.1101/2025.07.11.664293

**Authors:** David Patrick Kateete, Shakira Namakula, Edgar Kigozi, Fred A Katabazi, George William Kasule, Kenneth Musisi, Edward Wampande, Deus Lukoye, Moses L Joloba

**Author notes:** Corresponding authors: (DPK), (MLJ). These authors contributed equally to this work (David Patrick Kateete, Shakira Namakula, Edgar Kigozi).

## Abstract

Mutations in *rpoB*, a gene that encodes the bacterial RNA polymerase (RNAP) beta-subunit, can cause high-level resistance to rifampicin. Approximately 95% of rifampicin-resistant *Mycobacterium tuberculosis* clinical isolates possess mutations in an 81-base pair *rpoB* region referred to as the rifampicin-resistance determining region (*rpoB*/RRDR). Also, rifampicin-resistant *M. tuberculosis* clinical isolates carry multiple mutations in RNAP genes (i.e., *rpoA, rpoB, rpoC, rpoD*), particularly *rpoA* and *rpoC*, which encode the alpha-(α_2_) and beta′-(β′) subunits, respectively. Such secondary mutations offset the fitness cost associated with rifampicin-resistance mutations in *M. tuberculosis*, resulting in resistant strains that are as fit as the wildtype drug-susceptible strains. To analyse the patterns of compensatory mutations in RNAP encoding genes of rifampicin-resistant *M. tuberculosis* clinical isolates in Uganda, whole genome sequencing and Sanger DNA sequencing were performed on 52 *M. tuberculosis* clinical isolates – 20 drug-susceptible and 32 multidrug resistant (MDR). A total of 24 (75%) MDR-TB isolates had high-level rifampicin-resistance conferring mutations in *rpoB*/RRDR i.e., Ser531Leu (31%); His526Asp (6%); His526Leu (3%); His526Tyr (3%); His526Arg (3%); His526Gly (3%); Asp516Tyr (13%); Asp516Val (6%); Glu513Lys (3%); Leu511Pro (3%); Leu492Leu (3%); Gln490Arg (3%). Further, two putative compensatory mutations (Gln490Arg & Lys1025Glu) outside the RRDR and not resistance conferring were found in *rpoB*. Altogether, 16 (50%) MDR-TB isolates with *rpoB*/RRDR resistance conferring mutations had non-synonymous mutations in *rpoC* of the following patterns Leu39Phe (3%); Tyr61His (3%); Asp271Gly (3%); Ser377Ala (3%); Pro481Thr (3%); Val483Ala (6%); Leu516Pro (3%); Ala521Asp (3%); Gly594Glu (13%); Asn698Ser (3%); Leu823Pro (3%). In conclusion, putative compensatory mutations are prevalent in rifampicin-resistant *M. tuberculosis* clinical isolates in Uganda, with *rpoC*/Gly594Glu and *rpoC*/Val483Ala as the most frequent. Further studies will determine their association with strain genetic background, fitness and transmission in an endemic setting with a high burden of HIV-TB coinfection.

## Introduction

Multidrug resistance (MDR) in tuberculosis (TB), caused by *Mycobacterium tuberculosis* bacteria that are resistant to the two powerful anti-TB drugs (i.e., rifampicin and isoniazid [1, 2]), is a major threat to global TB control efforts and the attainment of the United Nations’ Sustainable Development Goal (SDG) #3 in the low- and middle-income countries (LMICs). Uganda is a TB endemic LMIC with high burden of TB and HIV coinfection, and relatively low MDR-TB rates i.e., ∼1.4% and ∼12.1% prevalence in newly diagnosed cases and retreated patients, respectively [3].

Bacterial RNA polymerase (RNAP) is an essential enzyme comprising five subunits i.e., α_2_ββ′ω that are responsible for DNA-dependent RNA synthesis [1, 4, 5]. The α_2_ββ′ω pentameric core forms a crab-claw-like structure where the alpha subunits (α_2_) are responsible for assembly while the ββ′ heterodimer forms the catalytic centre [6]. Unlike eukaryotic genomes that encode three distinct RNA polymerases, prokaryotes use one polymerase to synthesize rRNA, mRNA and tRNA, making the bacterial RNAP a fitness-determining enzyme as growth is dependent on the rate of production of rRNA [4, 6].

Rifampicin inhibits bacterial RNAP by binding to the beta (β) subunit, disrupting the DNA/RNA channel and blocking elongation of the RNA transcripts, which is bactericidal [1, 5, 6]. As such, mutations in *rpoB*, a gene that encodes the bacterial RNAP β-subunit, can cause high-level resistance to rifampicin; approximately 95% of *M. tuberculosis* clinical isolates resistant to rifampicin possess a non-synonymous mutation in an 81 base pair (bp) region of *rpoB* commonly referred to as the rifampicin-resistance determining region (RRDR; *rpoB*/RRDR), and such mutations often confer high-level resistance to rifampicin [1, 2, 7, 8]. Whilst antibiotic resistance in bacteria is associated with lower Darwinian fitness in the absence of antibiotics, clinical isolates of rifampicin-resistant bacteria do not suffer fitness deficit and often have levels of fitness similar to, or higher than those of wild-type drug-susceptible bacteria [1, 9]. (Here, ‘fitness’ refers to ‘the ability of a pathogenic bacterium to establish an infection, replicate and persist in an infected host, and its capacity to be efficiently transmitted’ to more susceptible hosts [10]). Furthermore, clinical isolates of rifampicin-resistant *M. tuberculosis* carry multiple mutations in the RNAP encoding genes (i.e., *rpoA, rpoB, rpoC, rpoD* [*rpoA/B/C/D*]) [1, 5], particularly *rpoA* and *rpoC* that encode the alpha (α_2_) and β′ subunits, respectively) [1, 5, 6]. It has been experimentally and/or epidemiologically demonstrated that these secondary mutations in *RNAP* genes offset the fitness cost associated with acquisition of rifampicin-resistance in *M. tuberculosis*, resulting in drug-resistant bacteria that are as fit as the wildtype drug-susceptible strains [1, 5, 6, 8, 9]. So far, it has been proven that compensatory mutations in *M. tuberculosis* (i) exhibit convergent evolution and commonly occur in rifampicin-resistant clinical isolates with mutations in the *rpoB/*RRDR, (ii) do not occur in rifampicin-susceptible clinical isolates nor in rifampicin-resistant clinical isolates without the *rpoB*/RRDR mutations [1, 6, 8-14] and, (iii) tend to be more prevalent in regions with the highest MDR-TB incidence in the world (e.g., South Africa and countries in Eastern Asia and/or Europe e.g., China, Russia, Abkhazia/Georgia, Uzbekistan, Kazakhstan, etc.) [1, 6, 9, 12-15].

As yet, little is known about the patterns of RNAP compensatory mutations among rifampicin-resistant *M. tuberculosis* clinical isolates from Uganda, a TB endemic country with a high-burden of both TB and HIV/AIDS. Here, we studied the *rpoA*/*B*/*C*/*D* gene sequences from clinical isolates of MDR- and pan-susceptible *M. tuberculosis* from patients in Uganda, for patterns of mutations associated with rifampicin-resistance. Characterization of such mutations contributes to our understanding of the microbiologic factors underlying antimicrobial resistance emergence, particularly the compensatory mechanisms allowing drug-resistant *M. tuberculosis* to overcome antibiotic pressure and become highly transmissible.

## Materials and Methods

### Study setting and isolates

This retrospective cross-sectional study was conducted on stored TB cultures at Makerere University College of Health Sciences in Kampala, Uganda, between 24^th^ June, 2019 and 17^th^ April, 2020. A total of 52 *M. tuberculosis* clinical isolates were studied, of which 20 were drug-susceptible and 32 were MDR-TB isolates i.e., rifampicin-resistant and isoniazid-resistant. The clinical and demographic characteristics of the patients, sputum sampling and TB culturing procedures were described previously in the national and Kampala drug-resistance surveys [3, 16]; briefly, the isolates were collected from a nationally representative cohort of new TB cases and previously treated sputum smear-positive patients registered at the TB diagnostic and treatment centres during 2009-2011 [3]. Similarly, phenotypic drug susceptibility testing for sensitivity to anti-TB drugs was described in the drug resistance surveys [3, 16] however, repeat culturing and susceptibility testing using the Löwenstein-Jensen (L-J) proportional method was done to validate the susceptibility patterns. *M. tuberculosis* culturing and drug sensitivity testing was performed in a BSL-3 Mycobacteriology Laboratory of the Dept. of Medical Microbiology, Makerere University School of Health Sciences.

### Extraction of chromosomal DNA

Stored bacterial isolates were accessed for sub-culturing on 1^st^ July, 2019; isolates were recovered by sub-culturing on Middlebrook 7H10 agar (Becton and Dickson, USA), incubating at 37ºC in a carbon dioxide incubator (Thermal Scientific, USA), and observing daily for 28 days for growth of the bacilli. The cells were harvested and suspended in absolute ethanol (Sigma scientific, USA) to kill them by suffocation. The suspension was centrifuged at 16,000 g to obtain the cell pellet that was later re-suspended in 0.25X Tris-EDTA (TE) buffer. High quality bacterial chromosomal DNA was extracted from *M. tuberculosis* by following the CTAB/chloroform extraction method [17]. Briefly, the cells, suspended in 0.25X TE buffer, were centrifuged at 3,000 g to wash off the media salts and residual ethanol. To lyse the cells, the cell pellet was re-suspended in 400 µL of fresh 0.25X TE buffer followed by adding 50 µL of lysozyme (40 mg/mL) and incubated at 37^º^C, overnight. Then, we added 150 µL of 10% Sodium Dodecyl Sulphate (SDS) / proteinase K solution to the mixture and incubated at 65^º^C for 1 hour to ascertain complete cell lysis, precipitation of proteins and other cell debris. After this, 100µL of 5M NaCl was added to each tube followed by 100 µL of CTAB/NaCl and inverting the mixture several times until the content turned milky. To ensure complete precipitation of all cell debris the solution was incubated briefly at 65^º^C for 10 minutes. To purify the DNA, 750 µL of chloroform/isoamyl alcohol mixture was added to the samples and centrifuged at 16,000 g for 10 minutes. After this, the aqueous phase that contained the DNA was carefully transferred to a pre-labelled sterile 1.5 ml microfuge tube, followed by adding 600 µL of absolute ice-cold isopropanol to precipitate the DNA. After incubating at -20^º^C for 2 hours, the mixture was centrifuged for 10 minutes at 16,000 g to pellet the DNA. The pellet was washed with 1 ml of 70% ice-cold ethanol and dried at room temperature for 1 hour. The DNA was eluted in 50 µL of 0.25X TE buffer. Prior to use in PCRs the quality and quantity of the extracted DNA was determined by electrophoresis on a 1% (w/v) agarose gel and a NanoDrop spectrophotometer (ThermoFisher Scientific).

### DNA sequencing and sequence analysis

The RNAP encoding genes (*rpoA, rpoB, rpoC, rpoD*) were PCR-amplified with previously published gene-specific primer sequences and procedures [7] for the *rpoB*/RRDR, and in-house primer sequences for *rpoA* (5’-CCGGTCACCATGTACCTACG-3’ forward, 5’-GGATGTCAAGCAGGTCGGAT-3’ reverse), *rpoC* (5’-CTACGTGATCACCTCGGTCG-3’ forward, 5’-GTTGACGATGATTTCCGGCG-3’ reverse), and *rpoD* (5’-CGATCGCGCGAAAAACCATCT-3’ forward, 5’-CACCGACTGCAGTTGATCCT-3’ reverse).

The targeted gene segments were successfully PCR-amplified from all the study isolates. The total reaction volume for all PCRs was 60 µL, prepared according to the HotStar PCR kit (QIAGEN, Hilden, Germany). Briefly, each reaction contained 27.5 µL nuclease free water, 6 µL 10x PCR buffer, 12 µL Q-solution, 3 µL 10 mM MgCl_2_, 3 µL of 10 mM dNTPs, 1.5 µL each of reverse and forward primers, 0.5 µL High-fidelity Taq DNA polymerase (5U/µL) (Sigma-Aldrich, USA), and 10 ng/µL (in 5µL volume) of the chromosomal DNA template. Amplification was achieved in a Thermocycler (Bio-Rad Laboratories Inc., Singapore) using the following program: initial denaturation at 95°C, 5 minutes, followed by 35 cycles each consisting of 95°C 45 seconds, 60°C 45 seconds, and 72°C 50 seconds, with a final extension step of 72°C, 10 minutes. Five microliters each of the PCR product was analysed using 1% agarose gel electrophoresis with Ethidium bromide 5mg/ml staining. Gels were run at 120 V for 1 hour and visualized using a UVP Gel documentation (Benchtop Trans-illuminator System_BioDoc-it, CA, USA). Then, 50 µL each of the PCR products was purified using the QIAmp DNA purification minikit (QIAGEN, Hilden, Germany), and the pure amplicons sequenced at ACGT Inc. (Wheeling IL, USA).

The sequences obtained were analysed first through BLAST searches at NCBI https://blast.ncbi.nlm.nih.gov/Blast.cgi to confirm they significantly align to the expected gene/protein sequences in *M. tuberculosis*. To determine the amino acid substitutions, the DNA sequences were translated into amino acid sequences using the Molecular Evolutionary Genetics Analysis Version 6.0 (MEGA6.06) software [7] or the ExPASy online database http://web.expasy.org/translate/. To identify mutations, the sequenced amplicons were aligned in MEGA6.06 and BioEdit v7.2.5.0 to the *rpoA, rpoB, rpoC and rpoD* gene sequences from *M. tuberculosis* H37Rv (NC_000962), a reference strain pan-susceptible to anti-TB drugs. To determine whether the identified mutations were resistance-conferring, we followed the tbdream database [18] https://tbdreamdb.ki.se/Info/ and published literature [1, 2, 6-9, 11, 13, 14]. All the data was curated, compiled and presented as tables or percentages depending on the frequency of mutations that occurred in the examined genes.

### Whole genome sequencing

Since some of the clinical isolates of rifampicin-resistant *M. tuberculosis* lacked rifampicin-resistance conferring mutations in the *rpoB*/RRDR upon analysis of the Sanger sequenced DNA amplicons, as a validation step, we performed whole genome sequencing (WGS) of all the 52 *M. tuberculosis* clinical isolates and the control strain H37Rv. WGS was performed at the Genomics Unit of the Department of Immunology and Molecular Biology at Makerere University College of Health Sciences, by following protocols published by illumina Inc. Briefly, 20 indexed paired-end libraries were prepared starting with 0.2 ng/µL of input *M. tuberculosis* chromosomal DNA using the Nextera® XT DNA sample preparation guide (illumina Inc., CA USA). This protocol included stages of tagmentation, indexing, normalization and library pooling prior to loading the pooled library into a MiSeq Reagent Kit v2 (MS-102-2003, 500 cycle) (illumina). This kit offers an improved chemistry to increase cluster density with a maximum output of 8.5 gigabyte (Gb) of data. The cartridge was loaded in the MiSeq instrument (illumina inc.) to run a resequencing workflow for ∼72 hours. The DNA library for WGS was spiked with PhiX v3 control (Illumina) to a final concentration of 5% recommended for low diversity libraries. The generated FASTQ files were analysed and high-quality files (Phred score of ≥30) uploaded on PhyResSE, a web-based tool that delineates *M. tuberculosis* antibiotic resistance markers and their lineages using WGS data (https://bioinf.fz-borstel.de/mchips/phyresse/).

### Quality control

PCR-amplicons from *M. tuberculosis* H37Rv, a reference strain that is susceptible to all anti-TB drugs, were sequenced and included as controls. Likewise, a known MDR-TB strain that contained well-known rifampicin and isoniazid resistance conferring mutations earlier confirmed by the Genotype MTBDR*plus* assay (Hain Life Sciences) was included in the study as the positive control for drug resistance mutations. Negative controls for DNA extraction, PCRs, and DNA sequencing were as well included to rule out contamination. High-Fidelity Taq DNA polymerase (Sigma-Aldrich, USA) that delivers superior results with twofold higher yield and threefold greater fidelity compared to regular Taq DNA Polymerase was used in the PCRs. Sanger DNA sequencing-identified mutations were revalidated through WGS.

### Ethics statement

This study was approved by the Makerere University School of Biomedical Sciences Research and Ethics Committee (approval # SBS-HDREC-667). This committee approved the use of archived *Mycobacterium tuberculosis* isolates previously collected from patients who participated in the National Drug Resistance Survey that investigated the phenotypic levels and patterns of resistance to first- and second-line anti-TB drugs among new and previously treated sputum smear-positive TB cases in Uganda [3, 16]. The parent studies stated herein had previously obtained written informed consent from the participants for sample storage and use of stored samples in further studies. The data were analysed anonymously and authors did not have access to information that could identify individual participants during or after data collection.

## Results and Discussion

We characterized a total of 52 *M. tuberculosis* clinical isolates; 32 MDR and 20 pan-susceptible to anti-TB drugs, for patterns of mutations in the *rpoA/B/C/D* genes associated with rifampicin-resistance in bacteria. The isolates were collected during the two drug resistance surveys conducted in the period between 2009 and 2011 [3, 16], and they are representative of the patients registered at the TB treatment centres at the time [3]. Overall, using both WGS and Sanger DNA sequencing, we did not find mutations or single nucleotide polymorphisms (SNPs) in *rpoA* and *rpoD* of all the isolates; likewise, there were no mutations/SNPs in any of the genes (*rpoA/B/C/D*) in the 20 drug-susceptible *M. tuberculosis* isolates and the control strain (H37Rv). As outlined below, mutations were found in the *rpoB* and *rpoC* genes in some of the MDR-TB isolates, and these were verified through WGS and Sanger DNA sequencing.

### *rpoB* mutations

**Fig 1** and **Table 1** summarize the patterns of mutations and SNPs we found in *rpoB*. A total of 24 MDR-TB isolates (75%, 24/32) had mutations in the *rpoB*/RRDR i.e., Lys1025Glu (3%, 1/32); Ser531Leu (31%, 10/32); His526Asp (6%, 2/32); His526Leu (3%, 1/32); His526Tyr (3%, 1/32); His526Arg (3%, 1/32); His526Gly (3%, 1/32); Asp516Tyr (13%, 4/32); Asp516Val (6%, 2/32); Glu513Lys (3%, 1/32); Leu511Pro (3%, 1/32); Leu492Leu (3%, 1/32); and Gln490Arg (3%, 1/32), and these are similar to mutations that have been reported before [2, 7, 10, 15] (note that three isolates had double mutations – see ahead). With the exception of Lys1025Glu, Leu492Leu and Gln490Arg, all mutations occurred in the *rpoB*/RRDR and confer high-level resistance to rifampicin [2, 8] apart from Leu511Pro, Asp516Tyr, and His526Gln that are categorized as low-level rifampicin-resistance conferring mutations [2, 8]. Therefore, apart from two mutations (i.e., Glu513Lys & Leu511P), rifampicin-resistance in this study is attributed to mutations at three notable codons – 516, 526, and 531, which is in line with reports from many settings around the world [2, 7]; notably, there is high polymorphism at codon 526 compared to the other two codons (516 & 531), **Fig 1**.

**Table 1:** Mutations in the *rpoB* and rpo*C* genes of the MDR-TB clinical isolates.

| Isolate # | *M. tuberculosis* genotype | *rpoB* (Rv0667) RRDR |  | *rpoC* (Rv0668) |  |
| --- | --- | --- | --- | --- | --- |
|  |  | Codon change | Mutation (*M. tuberculosis* coordinates) | Codon change | Mutation |
| 1. |  | TCG → TTG | Ser531Leu (S450L) | CTA → CCA | Leu516Pro |
| 2. | 4/Uganda-II | TCG → TTG | Ser531Leu (S450L) | CTA → CCA | Leu516Pro |
| 3. | 4/LAM | GAC → GTC | Asp516Val (D435V) | CTT → TTT | Leu39Phe |
|  |  |  |  | TAT → CAT | Tyr61His |
|  |  |  |  | GTG → GTT | Val68Val |
|  |  |  |  | TCT → GCT | Ser377Ala |
|  |  | AAA → GAA | Lys1025Glu (K944E) |  |  |
| 4. | 4/Uganda-II | GAC → TAC | Asp516Tyr (D435Y) | GGA → GAA | Gly594Glu |
| 5. | 4/LAM | GAC → TAC | Asp516Tyr (D435Y) | GGA → GAA | Gly594Glu |
| 6. | 4/Uganda-I | TCG → TTG | Ser531Leu (S450L) | GTA → GCA | Val483Ala |
| 7. | 4/Uganda-II | TCG → TTG | Ser531Leu (S450L) | GCT → GAT | Ala521Asp |
| 8. | 4/Uganda-I | CAA → AAA | Gln513Lys (Q432K) | CCT → ACT | Pro481Thr |
| 9. | 4/LAM | TCG → TTG | Ser531Leu (S450L) | AAC → AGC | Asn698Ser |
| 10. | 3/Delhi/CAS | CAC → CTC | His526Leu (H445L) | - |  |
| 11. | 4/Uganda-I | TCG → TTG | Ser531Leu (S450L) | GTA → GCA | Val483Ala |
| 12. | 4/LAM | CAC → TAC | His526Tyr (H445Y) | - |  |
| 13. | 4/LAM | CAC → GAC | His526Asp (H445D) | - |  |
| 14. | 4/EAM* | GAC → TAC | Asp516Tyr (D435Y) | GGA → GAA | Gly594Glu |
| 15. | 2/Beijing | TCG → TTG | Ser531Leu (S450L) | AGA → AGG | Arg84Arg |
| 16. | 3/Delhi/CAS | GTG → CCG | Leu511Pro (L430P) | GAC → GGC | Asp271Gly |
| 17. | 4/Uganda-I | CAC → CAA | His526Gln (H445Q) |  |  |
| 18. | 4/Uganda-II | CAC → CGC | His526Arg (H445R) | GCC → GCG | Ala542Ala |
| 19. | 4/LAM | CTC → CTT | Leu492Leu (L411L) | GTT → GTG | Val197Val |
|  |  | TCG → TTG | Ser531Leu (S450L) |  |  |
| 20. | 3/Delhi/CAS | CAA → CGA | Gln490Arg (Q409R) | CTT → CCT | Leu823Pro |
|  |  | TCG → TTG | Ser531Leu |  |  |
|  |  |  | (S450L) |  |  |
| 21. | 4/Uganda-II | TCG → TTG | Ser531Leu (S450L) | - |  |
| 22. | 3/Delhi/CAS | GAC → GTC | Asp516Val (D435V) | - |  |
| 23. | 3/Delhi/CAS | CAC → GAC | His526Asp (H445D) | - |  |
| 24. | 4/Uganda-I | GAT → TAT | Asp516Tyr (D435Y) | GGA → GAA | Gly594Glu |
| 25. | 4/Uganda-I | - |  | - |  |
| 26. | 2/Beijing | - |  | - |  |
| 27. | 4/LAM | - |  | - |  |
| 28. | 3/Delhi/CAS | - |  | - |  |
| 29. | 3/Delhi/CAS | - |  | - |  |
| 30. | 2/Beijing | - |  | - |  |
| 31. | 4/EAM* | - |  | - |  |
| 32. | 4/S | - |  | - |  |
|  | H37Rv (control) | - |  | - |  |

- No mutation detected; \*Euro-American super-lineage

**Fig 1.**
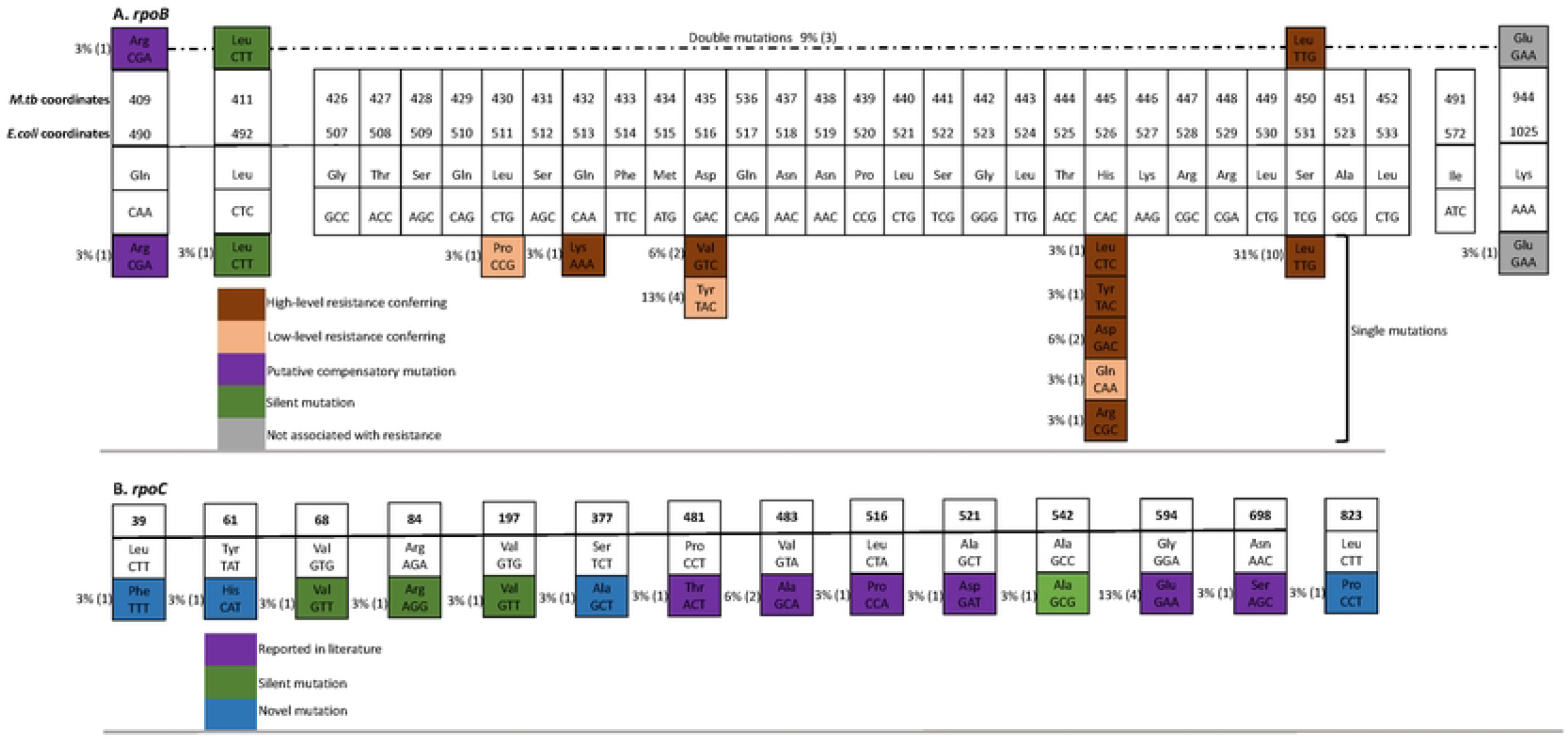
Rifampicin-resistance conferring mutations in *rpoB* (panel A) and putative compensatory mutations in *rpoB* (panel A) and *rpoC (*panel B) of MDR *M. tuberculosis* clinical isolates.

Furthermore, three isolates (#s 3, 19, 20) were double mutants (**Table 1, Fig 1**) in that each harboured two mutations i.e., Asp516Val & Lys1025Glu; Leu492Leu & Ser531Leu; and Gln490Arg & Ser 531Leu, respectively. Asp516Val and Ser531Leu are high-level rifampicin-resistance conferring mutations (see above) while the other three mutations in the double mutants occurred outside of the *rpoB*/RRDR and are not likely to be resistance conferring [2]; non-synonymous *rpoB* polymorphisms like Gln490Arg & Lys1025Glu that are outside of the *rpoB*/RRDR and are not resistance conferring have been reported to have compensatory effects [6] hence, are putative compensatory mutations.

### *rpoC* mutations and co-occurrence with mutations *rpoB*/RRDR

A total of 16 MDR-TB isolates (50%, 16/32) had mutations in *rpoC* – fifteen non-synonymous i.e., Leu39Phe (3%, 1/32); Tyr61His (3%, 1/32); Asp271Gly (3%, 1/32); Ser377Ala (3%, 1/32); Pro481Thr (3%, 1/32); Val483Ala (6%, 2/32); Leu516Pro (3%, 1/32); Ala521Asp (3%, 1/32); Gly594Glu (13%, 4/32); Asn698Ser (3%, 1/32); Leu823Pro (3%, 1/32) and four synonymous i.e., Val68Val (3%, 1/32); Arg84Arg (3%, 1/32); Val197Val (3%, 1/32); and Ala542Ala (3%, 1/32). Hence, the most frequent *rpoC* mutations were Gly594Glu (13%) and Val483Ala (6%), **Fig 1 & Table 1**. Of note, mutations Pro481Thr, Val483Ala, Leu516Pro, Ala521Asp, Gly594Glu and Asn698Ser are the most frequently reported *rpoC* compensatory mutations in rifampicin-resistant *M. tuberculosis* clinical isolates in many settings around the world [1, 6, 8-14]. While most of the mutations have been previously reported, we identified four new polymorphisms i.e., Leu39Phe, Tyr61His, Asp271Gly, and Ser377Ala, **Fig 1 & Table 1** that are likely to be compensatory mutations since the RNAP encoding genes are highly conserved in bacteria particularly *M. tuberculosis* [1, 9, 19]. Furthermore, 16 (67%) of the 24 MDR-TB isolates with rifampicin-resistance conferring mutations in *rpoB*/RRDR had non-synonymous mutations in *rpoC* (one had a synonymous mutation); in line with current evidence that compensatory mutations do occur only in rifampicin-resistant clinical isolates with rifampicin-resistance conferring mutations in *rpoB*/RRDR [1, 9, 12-14], we did not find mutations/SNPs in *rpoC* of the eight MDR-TB isolates that lacked rifampicin-resistance conferring mutations, **Table 1**.

Overall, in this study, the proportion of MDR-TB isolates with putative compensatory mutations in *rpoC* i.e., 50% (16/32) is high compared to rates from other countries [1, 8, 9, 12-14], and the fact that MDR-TB prevalence is low in Uganda [20, 21]; however, other investigators reported comparable or higher rates e.g., Wang et al in China [13] and Vargas et al in Peru [12] reported that 98.2% (54/55) and 54% (95/175) respectively, of rifampicin-resistant *M. tuberculosis* isolates with resistance-conferring mutations in *rpoB*/RRDR had putative compensatory nonsynonymous mutations in *rpoA*/*rpoC*.

Furthermore, almost all isolates (90%, 9/10) with the *rpoB*/Ser531Leu resistance conferring mutation harboured putative compensatory mutations in *rpoC* (Table 1), and similar observations for this mutation have been reported [8-11, 13, 14]. Furthermore, Gly594Glu, the most prevalent *rpoC* mutation in our study (**Table 1**) and reported before as a putative compensatory mutation [1, 6, 8-14], occurred among both rifampicin-resistant and rifampicin-susceptible isolates in Peru [12]. This discrepancy could be attributed to issues related to phenotypic drug susceptibility testing approaches i.e., the L-J proportional method used in the current study [3, 16] vs. slide drug susceptibility testing (SDST) in the Peruvian study [12]. Generally, phenotypic susceptibility testing methods, especially SDST, could yield false-negative results [22] for isolates with low-level resistance-conferring mutations leading to mis-identification of resistant isolates as susceptible. Furthermore, the synonymous mutation Ala542Ala was previously reported as a lineage-defining polymorphism (i.e., phylogenetic marker) for the Latin American Mediterranean (LAM) sub-lineage of the Euro-American *M. tuberculosis* super-family [9]; however, in this study, we did not detect Ala542Ala in LAM strains but instead was detected in *M. tuberculosis* sub-lineage Uganda, **Table 1**. This discrepancy could be related to the sensitivity of genotyping approaches used i.e., Spoligotyping/*IS6110* RFLP fingerprinting in the South African study [9], which is inferior and has a low discriminatory power compared to DNA sequencing used in the current study [23].

One limitation in our study is that we had fewer rifampicin-resistant TB isolates as Uganda as a country has low MDR-TB rates. However, findings from this study have implications for the understanding of fitness, transmissibility and antimicrobial resistance mechanisms in an endemic setting with high HIV-TB coinfection rates but low prevalence of MDR-TB.

## Conclusions

This study shows that putative compensatory mutations are prevalent in clinical isolates of rifampicin-resistant *M. tuberculosis* in Uganda with *rpoC*/Gly594Glu and *rpoC*/Val483Ala as the most frequent; also, for the first time, the study identifies putative compensatory mutations in *rpoB* (Gln490Arg and Lys1025Glu) of rifampicin-resistant *M. tuberculosis* in Uganda. Further studies are required to investigate the association of such mutations with the strain genetic background, as well as their effect on strain fitness and TB transmission in an endemic setting with high HIV-TB coinfection rates but low prevalence of MDR-TB.

## Acknowledgements

Research reported in this publication was supported by the Fogarty International Center of the National Institutes of Health under Award Number D43TW010319; the project was also supported in part by the EDCTP2 programme of the European Union (grant number TMA2018CDF-2357-MTI-Plus) to DPK. The content is solely the responsibility of the authors and does not necessarily represent the official views of the funders.

We thank the contribution and support given by staff at the Departments of Medical Microbiology and Immunology & Molecular Biology at the School of Biomedical Sciences, Makerere University College of Health Sciences in Kampala, Uganda.

## Notes

### Competing Interest Statement

The authors have declared no competing interest.

